# A zebrafish immunodeficiency model induced by the combination of Tacrolimus and Everolimus for rapid assessment of immune-enhancing agents

**DOI:** 10.1101/2025.04.06.647472

**Authors:** Mei Huang, Zhipeng Jia, Jun Zhou, Yiliang Liu, Xing Lu, Xiaozhen He

**Author notes:** To whom correspondence should be addressed. (Xiaozhen He); (Xing Lu). The authors wish it to be known that, in their opinion, the first two authors should be regarded as joint First Authors.

## Abstract

A methodology for the assessment of the immune-enhancing effects of various substances within a period of 5 days was reported. Experimentally, it was determined that the continuous treatment of Tacrollimus (0.5 ng/mL) and Everolimus (5 μg/mL) on zebrafish from 48 hpf to 120 hpf was sufficient to establish an immunodeficient model. The results showed that the model exhibited a significant reduction in the number of immune cells (thymocytes, myeloid cells, and macrophages) and a greater than 50% decrease in the expression of immune-related genes (*itk*, *hexb*, *cpa5*, *isg15*, *rsad2*). Astragalus powder can effectively increase the content of thymic T cells and macrophages, but its effect on myeloid cells is limited. Ganoderma lucidum spore powder had the most obvious influence on macrophages, with limited impact on thymus cells and myeloid cells. Royal jelly showed a significant effect on thymus T cells and macrophages, but its effect on myeloid cells was limited. Bailing capsule can significantly increase the content of myeloid cells and thymus T cells, while its effect on macrophages was limited. The study showed immune indexes varied in recovery with different immunomodulators versus the model group, suggesting they may act via distinct mechanisms.

**Summary Statement:** A robust immune system is vital to overall health and quality of life. Recent years have witnessed surging development of immune-boosting pharmaceuticals, nutraceuticals, and functional foods. However, an easy-to-use, rapid and intuitive immunity assessment framework is still missing. Here, a methodology for the assessment of the immune-enhancing effects of various substances within a period of 5 days was reported.

## Introduction

Long-term exposure to adverse environmental factors and certain lifestyle habits can impact the human immune system, potentially leading to immune dysfunction. As the demand for daily health care and immune boosting intensifies, particularly in light of the pneumonia epidemic’s influence, products alleged to enhance immunity have gained popularity. However, unlike medications prescribed for treating specific diseases, many substances purported to have immune-enhancing properties often lack rigorous biological validation. In recent years, there has been an uptick in misleading claims about immune-boosting substances, which poses risks to both the financial and physical well-being of consumers. Furthermore, the assessment of immune function has been hindered by the absence of more intuitive evaluation methods. Consequently, it is imperative to assess the immune-enhancing effects of substances with unknown efficacy, as well as those that are widely used, through appropriate models and indicators to ensure their safety and effectiveness.

As a model organism, zebrafish (*Danio rerio*) has the advantages of low cost, high throughput and visualization [1, 2]. Models of immunodeficiency in zebrafish can indeed be categorized into two main types, innate immunodeficiency and acquired immunodeficiency models. The innate immune system is active early in the zebrafish embryo, with macrophages produced in the cephalic mesenchyme before the onset of haematopoiesis (∼18 hpf) and myeloid cells already detectable at 48 hpf [3]. T cells and B cells of zebrafish begin to develop 4 days after fertilization (dpf), and functional maturity takes 4-6 weeks [4]. Zebrafish have key elements required for a recombinant activated gene (rag) - dependent adaptive immune system, including T cells, B cells, thymus, MHC antigens, and enzymes (such as terminal transferases) [5, 6]. Three different bacterial preparations were used to infect zebrafish, and the results showed that the temporal expression of acute phase proteins in zebrafish acute phase response was similar to that in humans, such as the rapid upregulation of pro-inflammatory cytokines IL-1β and TNFα, indicating that the acute phase response of humans to infection can be simulated through zebrafish models [6]. In order to achieve real-time visualization of the immune system for the detection and innate control, various zebrafish strains have been developed. For example, the promoters from *mpo* [7], *fli1* [8], and *lysC* [9] genes are used to drive fluorescence expression in myeloid cells, macrophages, macrophages, and granulocyte subpopulations, respectively. Zebrafish would be an ideal model organism for evaluating immune efficacy. Chemotherapeutic agents such as vinorelbine, chloramphenicol and immunosuppressive drug rapamycin are usually used to establish immunocompromised zebrafish models [10]. Typically, these models are established through immersion administration, although some studies may utilize intravenous injection. However, the existing zebrafish immune deficiency models are fraught with issues such as excessively high doses of modeling drugs, adverse side effects, and less than optimal outcomes. In addition, the injection method requires professional personnel and operation, and is not easy to perform high-throughput screening. Therefore, it is necessary to continuously optimize the zebrafish immunodeficiency model.

Immunosuppressants are a class of drugs that have been developed for organ transplantation to prevent rejection and autoimmune diseases, which can inhibit the immune response of the body. Tacrolimus (FK506), a class of cytokine inhibitor, is a macrolide immunosuppressant with potent immunosuppressive effects extracted from a soil fungus. FK506 inhibits lymphocyte proliferation, inhibits Ca^2+^ dependent activation of T-bar and B-lymphocytes, inhibits T-cell-dependent B-cells’ ability to produce immunoglobulins [11, 12]. Rapamycin is a macrolide isolated from the sap of *Streptomyces* hy-groscoppicus, and Everolimus is a derivative of immunosuppressive rapamycin, an mTOR inhibitor [13]. A phase III clinical trial using early combination of Everolimus and low-dose Tacrolimus was conducted in kidney transplant patients, confirming that the patients had better renal function, significantly reduced the incidence of acute rejection and graft failure, and the safety period could be extended to 12 months after transplantation [14]. It was found that controlling the blood concentration of Everolimus at 3-8 ng/mL, in combination with Tacrolimus, has good efficacy and safety in patients after heart transplantation, with an average survival rate of 95% at 3 years after surgery 74% [15]. The graft loss rate for the combination of Everolimus plus low-dose Tacrolimus is comparable to that of standard-dose Tacrolimus alone. However, the former regimen demonstrates significantly enhanced efficacy and is capable of improving renal function following transplantation [16]. Although the combination of immunosuppressants has a solid foundation for clinical application, it appears to be infrequently employed in the development of animal models for immunodeficiency.

In this study, the effects of Tacrolimus and Everolimus treatment, both individually and in combination, on the development and immunity function of zebrafish were comprehensively evaluated. Based on this, a new zebrafish immunodeficiency model has been established. Moreover, the corresponding immunity evaluation index was further optimized. Subsequently, the immunopotentiating effects of four different immune modulators (Astragalus powder, Royal jelly, Ganoderma lucidum spore powder and Bailing capsules) will be evaluated. These findings provide a good theoretical basis for the efficacy evaluation and development of immunomodulators.

## Results

### Acquisition of immunodeficiency model

We hypothesized that the combined treatment of Tacrolimus and Everolimus could establish a highly efficient model of immunodeficiency in zebrafish, which could complete the efficacy evaluation of candidate immunomodulators within 5 days (Fig. 1A). We conducted preliminary experiments to determine the concentrations of Tacrolimus and Everolimus, both individually and in combination. The combination of preliminary group 1 and preliminary group 2, which used lower drug concentrations, did not show a significant difference in the malformation rate and mortality rate of developing embryos at 1-5 dpf when compared to the control group. In contrast, the combination preliminary group 3 and preliminary group 4 which involved higher drug concentrations, exceeded the predefined selection criteria, with mortality rate not exceeding 30% (Supplemental Table 1 and Supplementary Fig. 1A).

**Fig. 1.**
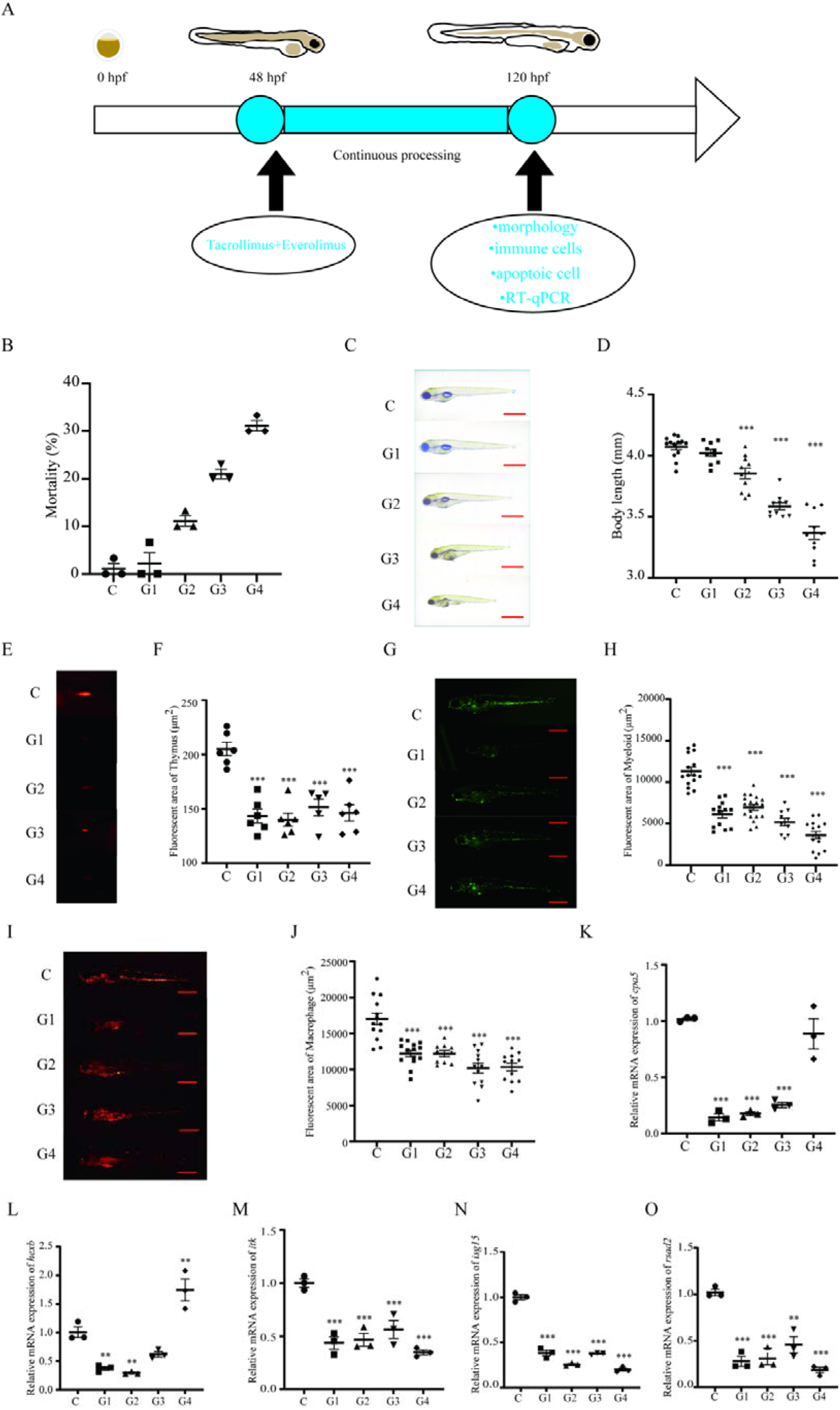
Acquisition of immunodeficiency model. (A) Experiments described in the timeline. (B) Mortality rate of Control group, G1, G2, G3 and G4 120 hpf zebrafish. (C) Control group, G1, G2, G3 and G4 120 hpf zebrafish pictures with a scale of 1 mm. (D) Control group, G1, G2, G3 and G4 120 hpf zebrafish body length statistics. (E and F) Control group, G1, G2, G3 and G4 120 hpf zebrafish thymus pictures (E) and thymus fluorescence area statistics(F). (G and H) Control group, G1, G2, G3 and G4 120 hpf zebrafish myeloid cell pictures (G) and myeloid cell fluorescence area statistics(H). (I and J) Control group, G1, G2, G3 and G4 120 hpf zebrafish macrophage pictures (I) and macrophage fluorescence area statistics (J). (K and O) Immune-related gene expression in Control group, G1, G2, G3 and G4 120 hpf zebrafish. All data were compared using one-way ANOVA, and *P*-values reflect differences between experimental groups,*=*p*<0.05; **=*P*<0.01; ***=*P*<0.001vs. Control group.

The preliminary group 3 group exhibited a mortality rate of 47%, slightly exceeding the acceptable threshold. Therefore, the combination of 3.5 ng/mL Tacrolimus and 20 μg/mL Everolimus, representing the highest concentrations, was selected for use in the formal experimental group. Concurrently, the lowest concentration used in the preliminary experiment of G1 group did not affect the embryonic phenotype (Supplemental Table 2). To optimize the immunosuppressive effect, the formal experimental group will employ the lowest effective concentration, which is 0.5 ng/mL of Tacrolimus combined with 5 μg/mL of Everolimus. Thus, the formal experimental group will set the minimum concentration at 0.5 ng/mL of Tacrolimus in conjunction with 5 μg/mL of Everolimus. To investigate whether the timing of treatment initiation at 24 hpf or 48 hpf influenced the outcomes of model establishment, the findings revealed no significant disparities in mortality rates between the two treatment timelines (Supplementary Fig. 1B). This suggests that the developmental toxicity profile of zebrafish was comparable for both early and later treatment initiation times. Meanwhile, considering that zebrafish embryos may undergo spontaneous death from fertilization to hatching due to quality issues, in order to avoid errors in the experiment caused by this situation, drug treatment was chosen to start at 48 hpf.

We administered varying concentrations of experimental drug on zebrafish between 48-120 hpf, recorded the phenotypic outcomes, and assessed the toxicity profile of different drug mixtures on the zebrafish larvae. Across the experimental groups, a trend emerged where an increase in drug concentration corresponded with a rise in the incidence of mortality rate, starting from less than 10% at the lowest concentration in G1 to over 30% at the highest concentration in G4 (Fig. 1B). At higher concentrations, malformations such as spinal curvature, pericardial and yolk sac enlargement, and intestinal defects were markedly more pronounced (Fig. 1C). Moreover, a significant reduction in body length was noted with escalating drug concentrations (Fig. 1D). The findings suggest that the combination of Tacrolimus and Everolimus exerts developmental toxicity on zebrafish embryos, resulting in malformations at higher concentrations. However, at lower concentrations, the phenotypes did not significantly differ from those of the control group, leading us to propose that the developmental toxicity of combined immunosuppressant therapy on zebrafish embryos is reduced at subtoxic concentrations.

In order to investigate the concentration ratio that would most effectively inhibit the immune system, we treated T cells, myeloid cells and macrophages transgenic zebrafish with various drug concentrations and assessed the impact on immune cell populations. At 120 hpf, we photographed and quantified the fluorescence area as an indicator of immune cell abundance. The results revealed a significant decrease in the fluorescence area of the three types of immune cells in the experimental groups when compared to the control group (Fig. 1E-I). Notably, the thymus area exhibited the most pronounced reduction, with G1, G2, and G4 showing nearly a 50% decrease (Fig. 1E, F).The fluorescence area of myeloid cells significantly decreased following immunosuppressant treatment, with considerable differences observed among the treatment groups (Fig. 1G, H). G2 demonstrated the least inhibition, with approximately a 35% reduction, while G4 had the most significant inhibitory effect, with a reduction exceeding 50%. Similarly, The inhibitory effect on macrophages across the four groups showed less variation, with the fluorescence area ranging from 30% to 40% (Fig. 1I, J). In summary, the four concentration gradient treatments showed varying degrees of inhibition on the fluorescence of the three immune cell types, but overall, the differences in inhibition among the groups were not pronounced.

To ascertain whether the compromised immunity and reduction in immune cells in zebrafish, induced by the drug combination, were a result of enhanced apoptosis, we conducted AO staining to visualize and quantify the number of apoptotic cells in the treated zebrafish (Supplementary Fig. 1E, F). Contrary to expectations, the number of apoptotic cells in the experimental group did not rise but actually decreased in comparison to the control group. Moreover, there was no significant difference observed in the number of apoptotic light spots among the various concentration-treated groups (Fig. 1K, O). We specifically examined the tail region, where the most pronounced decline in immune cells was noted. Despite the reduction in immune cell numbers, we did not observe an increase in apoptosis within the tail. This finding suggests that the suppression of immune cell numbers by the drug treatment is not mediated through apoptosis. Instead, it may be achieved by inhibiting cell production or other mechanisms that do not involve programmed cell death.

### Combination of Tacrolimus and Everolimus gives better immunodeficiency results than alone

To further determine the immunosuppressive effect of the combination of Tacrolimus (0.5 ng/mL) and Everolimus (5 μg/mL) is more potent than the effect of each drug administered individually. We treated zebrafish embryos with the combination alone. We then observe the fluorescence area of immune cells to determine if there was a more pronounced decrease when compared to the combination treatment. The statistical results revealed that the fluorescence areas of the thymus, myeloid cells, and macrophages in the groups treated with Tacrolimus or Everolimus alone were significantly greater than those in the drug combination group (Fig. 2A-F). This indicates that the combination exerts a significantly stronger immunosuppressive effect than either drug alone. Consequently, we concluded that the treatment regimen of 48-120 hpf with the combination is optimal for establishing an immunodeficiency model in zebrafish.

**Fig. 2.**
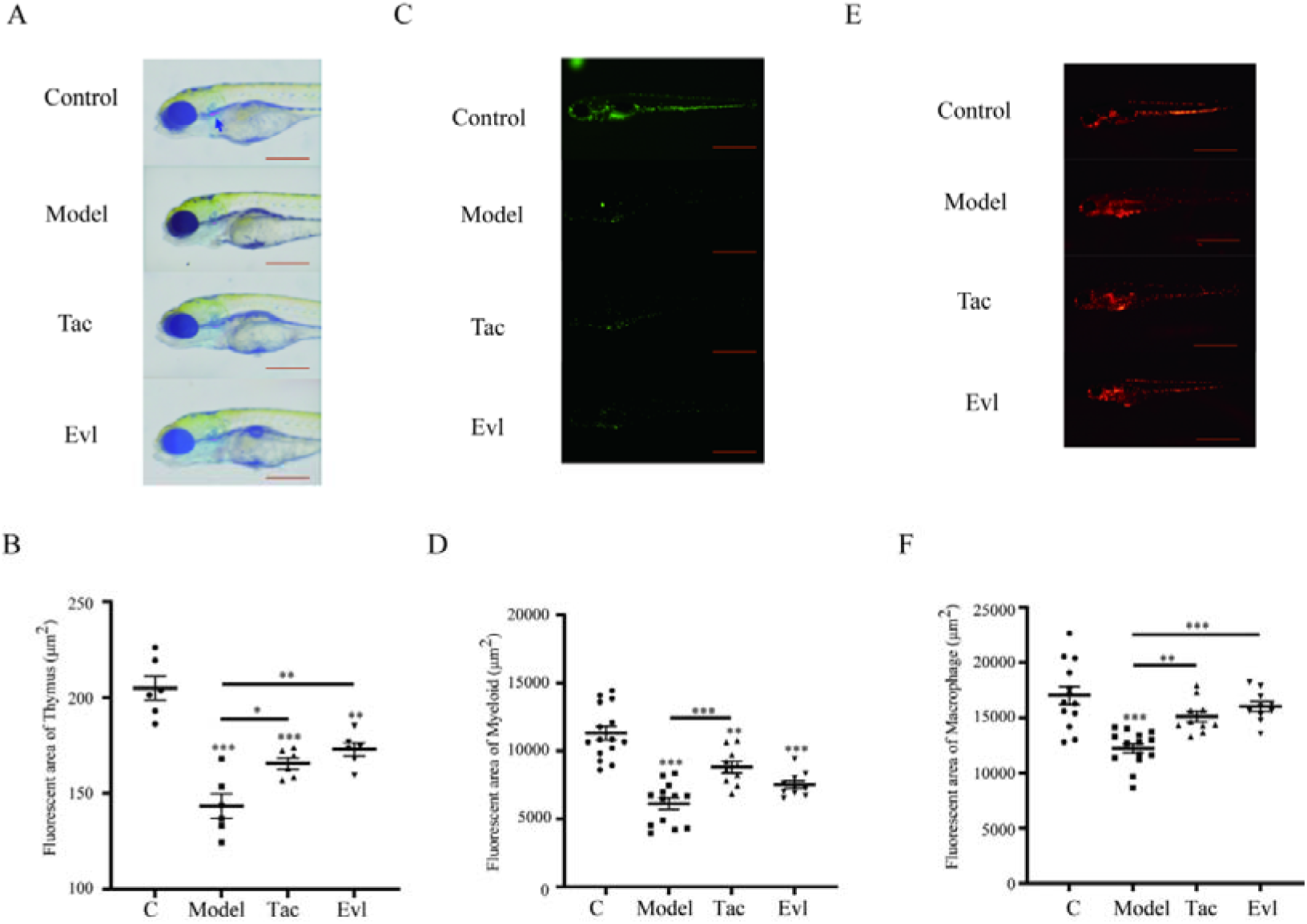
Combination of Tacrolimus and Everolimus gives better immunodeficiency results than alone. (A and B) Control group, thymus fluorescence (left) and area statistics (right) at 120 hpf in the immunodeficiency model group, Tacrolimus alone treatment and Everolimus alone treatment. (C and D) Control group, myeloid cell fluorescence (left) and area statistics (right) at 120 hpf in the immunodeficiency model group, Tacrolimus alone treatment and Everolimus alone treatment. (E and F) Control group, macrophage fluorescence (left) and area statistics (right) at 120 hpf in the immunodeficiency model group, Tacrolimus alone treatment and Everolimus alone treatment. Data were compared using one-way ANOVA, and *P*-values reflect differences between experimental groups, *=*p*<0.05; **=*p*<0.01; ***=*p*<0.001vs. Control group. Data were analyzed using the t-test and are presented as the mean values from three replicate experiments. Significance levels are indicated as follows: * *p* < 0.05, ** *p* < 0.01, *** *p* < 0.001.

### Immunomodulator-Astragalus powder evaluation results

To further validate the utility of our constructed immunodeficiency model for assessing immune function, we selected Astragalus powder, a potential immune enhancer, for evaluating its effects on enhancing immunity. The concentrations of the immunomodulators used were determined in a pre-experiment (Supplementary Table S3). Treatment with Astragalus powder commenced at 48 hpf and continued until 120 hpf in rows of immunocompetent zebrafish. The effects of Astragalus powder on the zebrafish were found to be concentration-dependent, with increased immune enhancement observed as the drug concentration increased (Supplementary Fig. 2A). To establish a range of concentrations for the treatment, we selected the highest drug concentration that resulted in a zebrafish mortality rate of less than 40%. Based on this, we set up three concentration levels in an ascending hierarchy for treating the modeling group (Supplementary Table S4).

We quantified the involvement of cells in immunoregulation and the expression of immunoregulatory genes as indicators. Thymic T-cell area in the group treated with Astragalus powder showed an increase relative to the model group, with the 1 mg/mL concentration exhibiting the most significant enhancement. However, the increase in myeloid cell content was modest, whereas a substantial restorative impact on macrophages was observed (Fig. 3A-F). The enhanced effect of Astragalus powder on macrophages may be attributed to their participation in a diverse array of physiological activities and multiple gene pathways, rendering them more susceptible to immunomodulatory influences. In terms of the expression of immunomodulatory genes, *cpa5* gene was down-regulated by up to 90% in the model group, and all concentrations of Astragalus powder demonstrated some degree of restorative effect, with 1 mg/mL proving to be the most effective (Fig. 3G). A significant up-regulation of the *hexb* gene was observed at a 1 mg/mL concentration of Astragalus powder compared to the model group (Fig. 3H). The expression of the *isg15* gene increased with higher concentrations of Astragalus powder, indicating a positive correlation with the treatment dose (Fig. 3I). No notable impact on the *itk* gene expression was detected following treatment with Astragalus powder (Fig. 3J). A slight up-regulation of the *rsad2* gene was seen in the Astragalus powder-treated group relative to the model group (Fig. 3K). In summary, the immunomodulator Astragalus powder exerts a degree of immunomodulatory influence on the immunodeficiency model. This also suggests that the immunodeficiency model we developed is suitable for immune function evaluation.

**Fig. 3.**
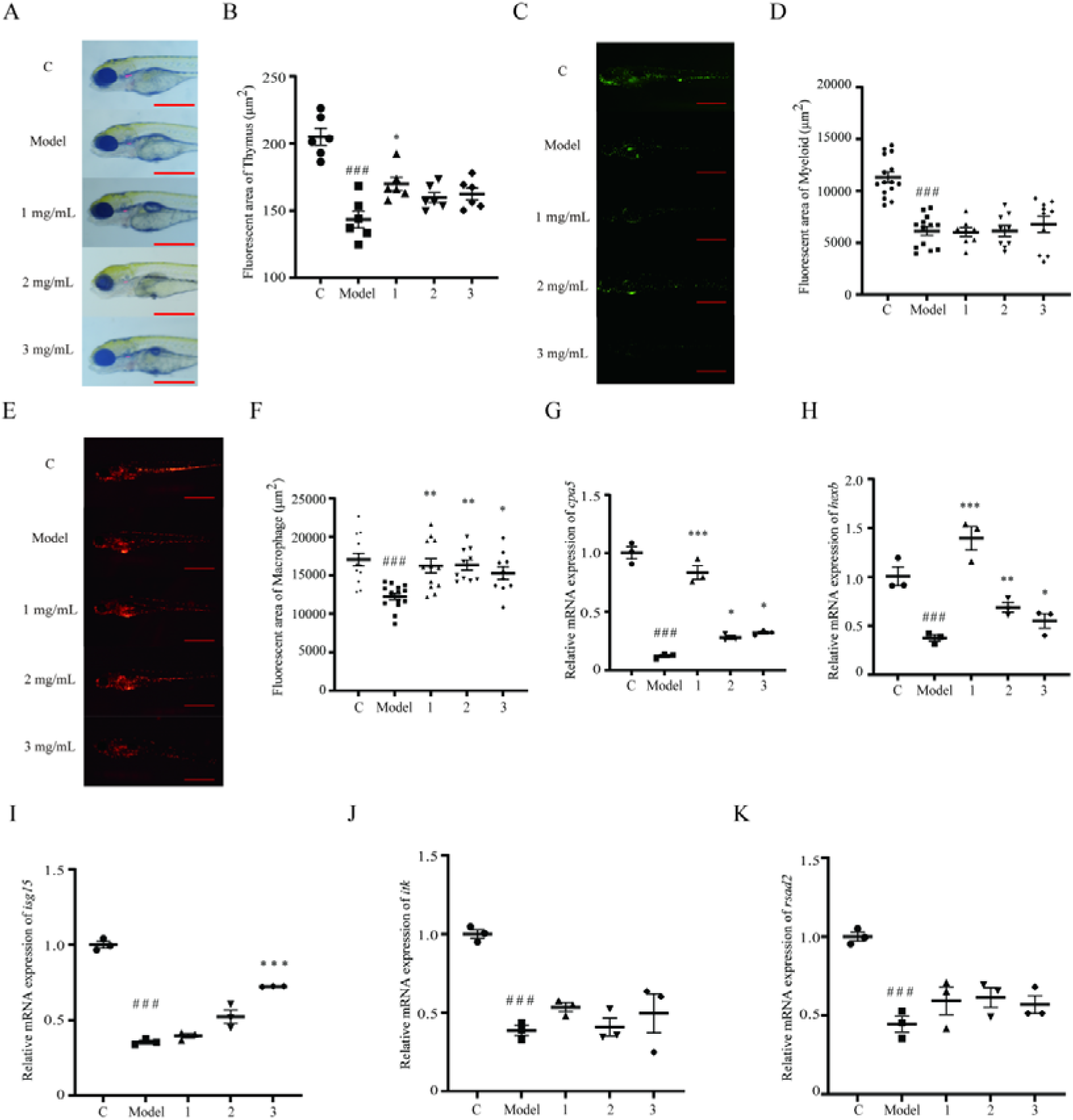
Immunomodulator-Astragalus powder evaluation results. (A and B) Thymus pictures and thymus fluorescence area statistics of Control group, model group, and the three groups with 1, 2, and 3 mg/mL Astragalus powder added at the same time of modelling. (C and D) myeloid cell pictures and myeloid cell fluorescence area statistics of Control group, model group, and the three groups with 1, 2, and 3 mg/mL Astragalus powder added at the same time of modelling. (E and F) Macrophage pictures and macrophage fluorescence area statistics of the Control group, the model group, and the three groups in which 1, 2, and 3 mg/mL Astragalus powder was added at the same time of modelling. (G and K) Immune-related gene expression in Control group, the model group, and the three groups in which 1, 2, and 3 mg/mL Astragalus powder was added at the same time of modelling. Data were compared using one-way ANOVA, and *P*-values reflect differences between experimental groups, *=*p*<0.05; **=*p*<0.01; ***=*p*<0.001vs. Model group. Data were analyzed using the t-test and are presented as the mean values from three replicate experiments. Significance levels are indicated as follows: # *p* < 0.05, ##*p* < 0.01, ### *p* < 0.001vs. Control group.

### Immunomodulator-Royal Jelly Evaluation Results

Our findings indicate that the treatment group receiving 200 μg/mL of Royal jelly demonstrated a substantial enhancement in the T-cell population within the thymus, which was beneficial but did not significantly impact the myeloid cell count. Additionally, the area occupied by macrophages in the 100 and 200 μg/mL concentration groups significantly diverged from that of the model group and closely approximated the control group. However, treatment at a concentration of 300 μg/mL showed a reduced efficacy in promoting thymus and macrophage recovery compared to the 100 and 200 μg/mL doses (Fig. 4A-F). The immunomodulatory effects of Royal jelly were evident in the up-regulation of the *cap5* gene when compared to the model group, with the 200 μg/mL Royal jelly group showing no significant difference in *cap5* gene expression from the control group (Fig. 4G). All tested concentrations of Royal jelly exhibited some degree of up-regulation for the *hexb* gene (Fig. 4H). Royal jelly at a concentration of 100 μg/mL showed moderate up-regulation of the *isg15* gene, though this was not statistically significant, whereas the 200 μg/mL and 300 μg/mL concentrations displayed abnormally high up-regulation (Fig. 4I). Royal jelly’s most notable impact was on the restoration of *itk* expression, with significant enhancements observed at 200 and 300 μg/mL compared to the model group (Fig. 4J). Concentrations of 200 μg/mL and 300 μg/mL of Royal jelly significantly up-regulated the *rsad2* gene compared to the model group, bringing it to levels nearly equivalent to those of the control group (Fig. 4K). These results imply that the immunodeficiency model we developed is suitable for assessing the immunomodulatory properties of Royal jelly.

**Fig. 4.**
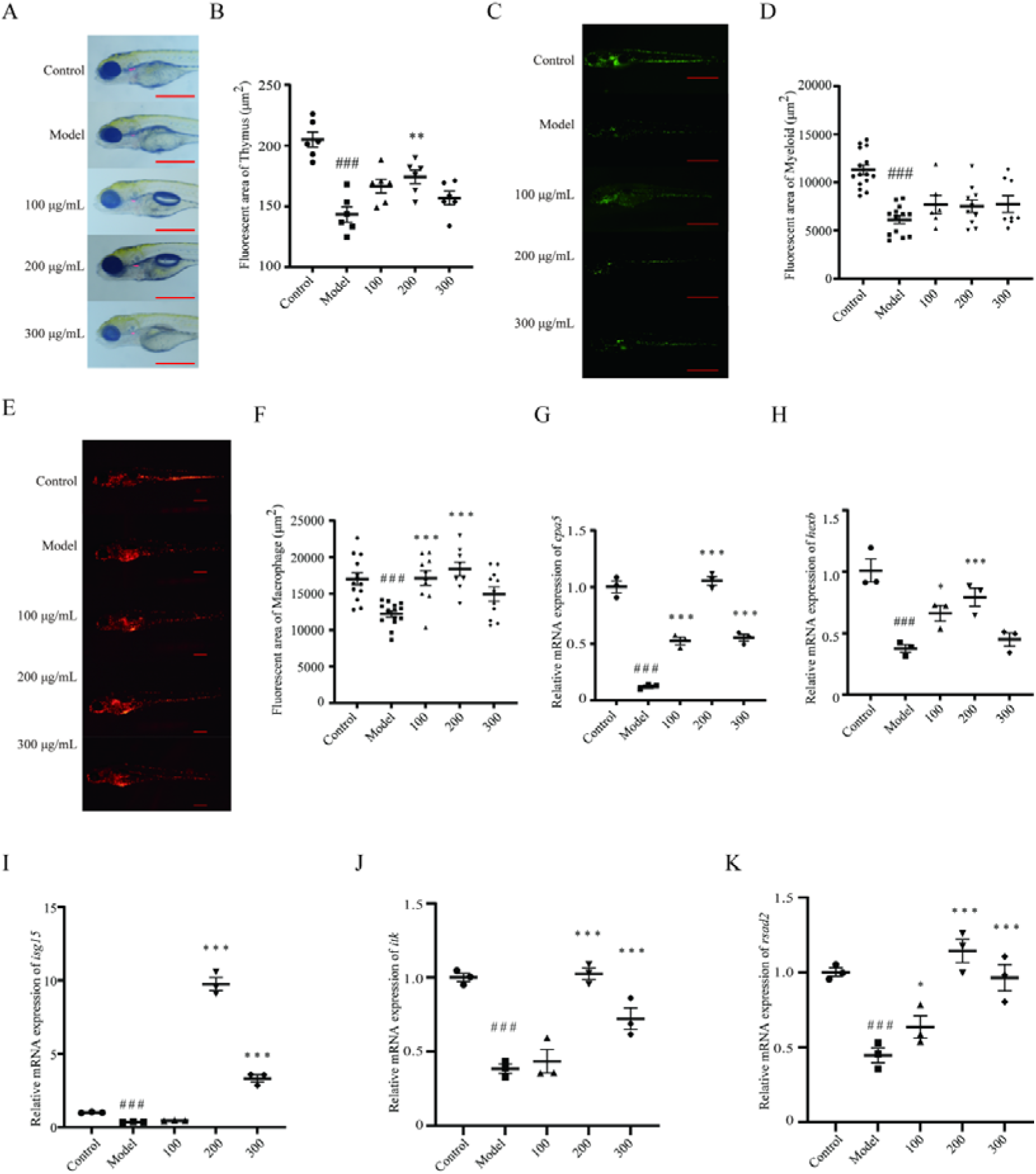
Immunomodulator-Royal Jelly Evaluation Results. (A and B) Thymus pictures and thymus fluorescence area statistics of Control group, model group, and three groups in which 100, 200, and 300 μg/mL Royal jelly was added at the same time of modelling. (C and D)Myeloid cell pictures and medullary cell fluorescence area statistics of Control group, model group, and three groups in which 100, 200, and 300 μg/mL Royal jelly was added at the same time of modelling. (E and F) Macrophage pictures and macrophage fluorescence area statistics of Control group, model group, and three groups with 100, 200, and 300 μg/mL Royal jelly added at the same time of modelling (G and K)Immune-related gene expression in Control group, model group, and three groups with 100, 200, and 300 μg/mL Royal jelly added at the same time of modelling. Data were compared using one-way ANOVA, and *P*-values reflect differences between experimental groups, *=*p*<0.05; **=*p*<0.01; ***=*p*< 0.001vs. Model group. Data were analyzed using the t-test and are presented as the mean values from three replicate experiments. Significance levels are indicated as follows: # *p* < 0.05, ##*p* < 0.01, ### *p* < 0.001vs. Control group.

### Immunomodulator-Ganoderma lucidum spore powder Evaluation Results

The Ganoderma lucidum spore powder showed limited ability to enhance the thymic T-cell content (Fig. 5A, B). There was some increase in myeloid cell count, but the results were not statistically significant (Fig. 5C, D). All three concentrations of the wall-broken Ganoderma lucidum spore powder (100, 500, and 1000 μg/mL) significantly increased the content of macrophages (Fig. 5E, F). The wall-broken Ganoderma lucidum spore powder up-regulated the expression of the *cap5* gene compared to the model group, with the most significant up-regulation observed at a concentration of 500 μg/mL (Fig. 5G). The *hexb* gene expression was also up-regulated by the spore powder, with the expression at 500 μg/mL and 1000 μg/mL exceeding that of the control group (Fig. 5H). At a concentration of 1000 μg/mL, the *isg15* gene showed significant up-regulation compared to the model group, reaching levels close to the control group (Fig. 5I). The wall-broken Ganoderma lucidum spore powder did not significantly restore the down-regulation of the *itk* gene (Fig. 5J). Similar to *itk*, the *rsad2* gene expression did not show significant restoration (Fig. 5K). These results suggest that the spore breaking powder of Ganoderma lucidum has various immunomodulatory effects, particularly on macrophage content and certain gene expressions, but may not be as effective in restoring T-cell content or specific gene expressions to control levels.

**Fig. 5.**
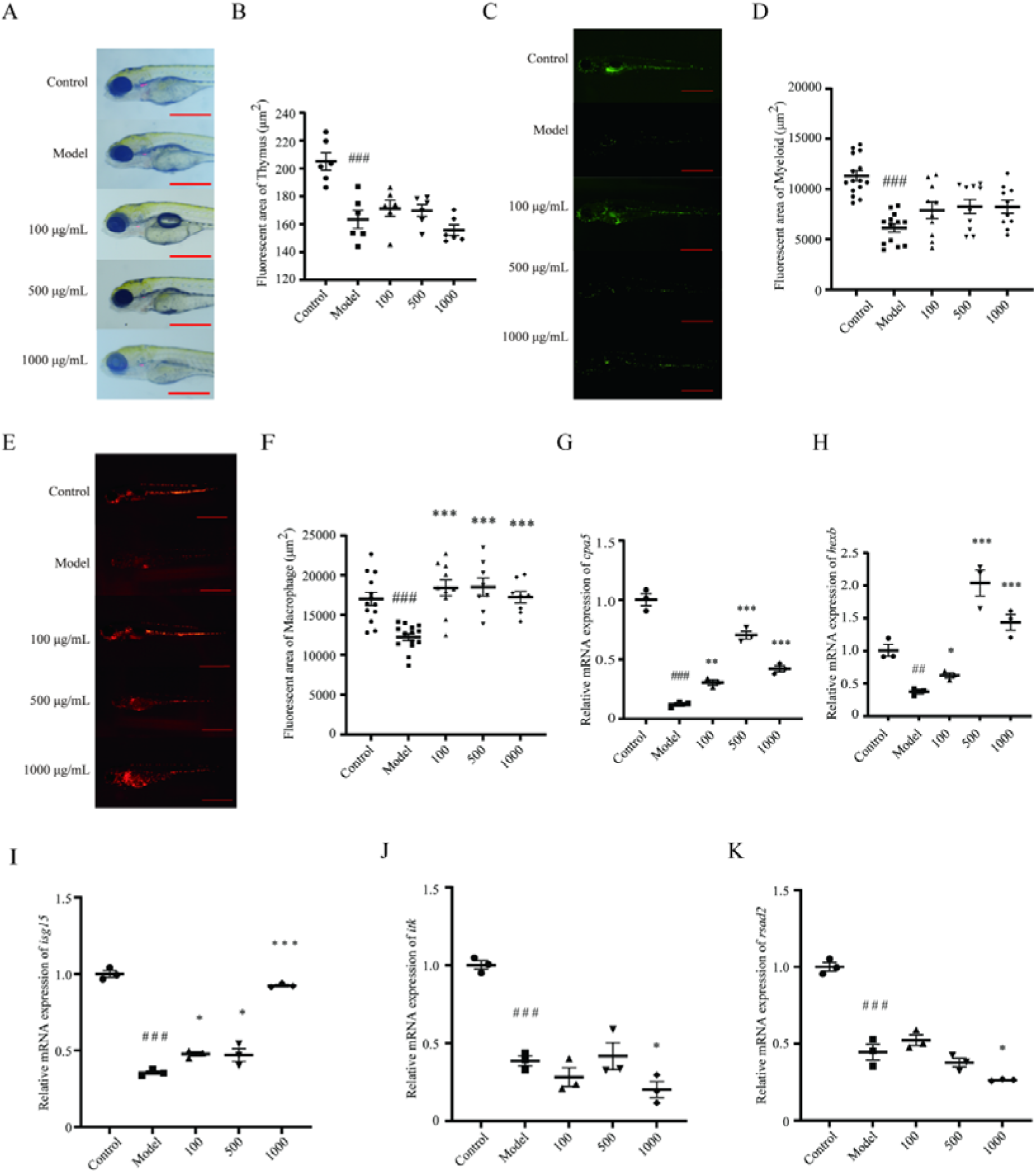
Immunomodulator-Ganoderma lucidum spore powder Evaluation Results. (A and B) Thymus pictures and thymus fluorescence area statistics of Control group, model group, and three groups in which 100, 500, and 1000 μg/mL Ganoderma lucidum spore powder was added at the same time of modelling. (C and D) Myeloid cell pictures and myeloid cell fluorescence area statistics of Control group, model group, and three groups in which 100, 500, and 1000 μg/mL Ganoderma lucidum spore powder was added at the same time of modelling. (E and F) Macrophage pictures and macrophage fluorescence area statistics of Control group, model group, and three groups in which 100, 500, and 1000 μg/mL Ganoderma lucidum spore powder was added at the same time of modelling. (G and K) Immune-related gene expression in Control group, model group, and three groups in which 100, 500, and 1000 μg/mL Ganoderma lucidum spore powder was added at the same time of modelling.Statistical significance analyses were calculated by Unpaired T-tests analysis. Data were compared using one-way ANOVA, and *P*-values reflect differences between experimental groups, *=*p*<0.05; **=*p*<0.01; ***=*p*< 0.001vs. Model group. Data were analyzed using the t-test and are presented as the mean values from three replicate experiments. Significance levels are indicated as follows: # *p* < 0.05, ##*p* < 0.01, ### *p* < 0.001vs. Control group.

### Immunomodulators - Bailing Capsules Evaluation Results

Bailing at 20 μg/mL significantly increased the fluorescence area of thymic T cells, an effect that diminished as the drug concentration rose (Fig. 6A, B). It also markedly elevated myeloid cell levels (Fig. 6C, D). However, its capacity to restore macrophages was minimal, with a notable decrease in the fluorescence area of macrophages even at high concentrations (Fig. 6E, F). Both 50 μg/mL Bailing treatments significantly up-regulated the *cpa5* gene (Fig. 6G). Bailing at 50 μg/mL also showed a significant up-regulation of *hexb* compared to the model group (Fig. 6H). The regulation of *isg15* by Bailing exhibited an inverse relationship with concentration, demonstrating a positive restorative effect at lower concentrations and a decline in regulation at higher concentrations (Fig. 6I). Bailing at 20 μg/mL significantly up-regulated *itk* (Fig. 6J), but there was no significant restoration of *itk* down-regulation by Bailing (Fig. 5K).

**Fig. 6.**
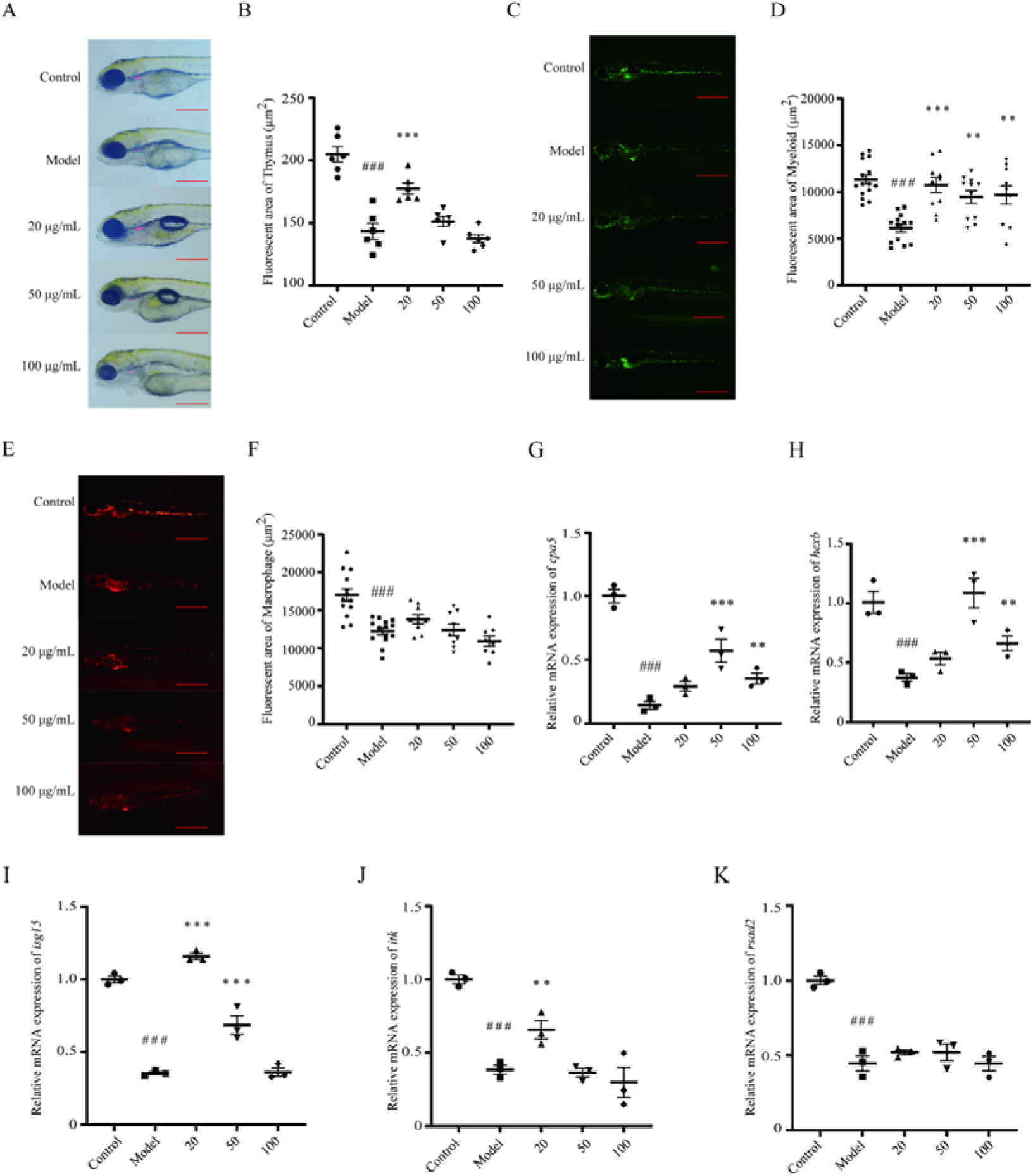
Immunomodulators - Bailing Capsules Evaluation Results. (A and B) Thymus pictures and thymus fluorescence area statistics for the Control group, the model group, and the three groups in which 20, 50, and 100 μg/mL of Bailin capsules were added at the same time as modelling. (C and D) Statistics of myeloid cell pictures and myeloid fluorescence area in Control group, model group, and three groups with 20, 50, and 100 μg/mL of Bailing capsule added at the same time of modelling. Statistical significance analyses were calculated by Unpaired T-tests analysis. (E and F) Macrophage pictures and macrophage fluorescence area statistics for the Control group, the model group, and the three groups in which 20, 50, and 100 μg/mL of Bailin capsules were added at the same time as modelling. (G and K) Immune-related gene expression in Control group, model group, and three groups in in which 20, 50, and 100 μg/mL of Bailin capsules were added at the same time as modelling. Data were compared using one-way ANOVA, and *P*-values reflect differences between experimental groups, *=*p*<0.05; **=*p*<0.01; ***=*p*<0.001vs. Model group. Data were analyzed using the t-test and are presented as the mean values from three replicate experiments. Significance levels are indicated as follows: # *p* < 0.05, ##*p* < 0.01, ### *p* < 0.001vs. Control group.

## Discussion

The combination treatment of Tacrolimus and Everolimus resulted in significant changes in the phenotype of zebrafish juveniles, evident through the development of spinal curvature, pericardial swelling, yolk sac swelling, intestinal malformations, as well as a decrease in myeloid cells, thymic T cells, and macrophages. Genes associated with immune system were either downregulated or upregulated, effectively establishing a zebrafish immunodeficiency model that rapidly caused immune defects in zebrafish at lower concentrations. In addition, we integrated the treatment with immune modulators and conducted statistical analyses to evaluate the regulatory effect of this model on immune-related indicators. The model has been shown to quickly assess certain immune regulatory substances in a preliminary manner. Pharmacological modeling offers benefits such as easy operation, reduced time requirements, and high throughput capabilities. Tacrolimus exerts its immunosuppressive effect *in vivo* by binding to the specific protein target, FK binding protein 12 (FKBP12), which is member of the immunophilin family. The complex that arises from the binding of Tacrolimus to FKBP12 interacts with calcineurin and inhibits its function by blocking the dephosphorylation of the nuclear factor of activated T cells (NFAT), preventing NFAT from translocating into the nucleus of activated T cells. This process suppresses the transcription and synthesis of downstream cytokines, such as interleukin-2 (IL-2), a potent stimulator of T cell activation and proliferation, and tumor necrosis factor alpha (TNF - α), which is associated with inflammatory responses, and interferon gamma (IFN-γ), among others [17]. Mechanistic target of rapamycin (mTOR) is a ser/thr kinase that, as a key signal transduction protein, has an inhibitory effect on antigen-induced T or B cell proliferation in cells. Rapamycin is a selective inhibitor of mTOR that exhibits poor solubility in water but can be dissolved in various organic solvents. This compound is a derivative isolated from the culture medium of *Streptococcus suis* fermentation and is classified as a macrolide antibiotic. The mode of action for rapamycin involves its association with the cytoplasmic protein-FK506 binding protein 12 (FKBPl2) in the cytoplasmic matrix, leading to thw formation of the rapamycin-FKBPl2 complex. This complex then interacts with mTOR, resulting in the inhibition of mTOR complex 1 (mTORCl) and affects gene transcription and various metabolic pathways. Everolimus, known by its developmental code RAD001, is a derivative of rapamycin and represents a novel class of mTOR inhibitors [18]. Our findings reveal that the combined administration of Tacrolimus and Everolimus has a significantly stronger immunosuppressive effect compared to when each drug is used independently (Fig. 2). We quantified the numbers of immune cells in zebrafish embryos that treated with Tacrolimus and Everolimus individually and/or in combination, and found that the drug combination group showed a significant increase in the number of thymus, myeloid cells, and macrophages when compared to the treatment alone.

Zebrafish *Tg (lyz: DsRED2)* it is a macrophage reporter line in which the zebrafish lysC promoter was used to drive macrophage expression of DsRED2[9]. The GFP fluorescence signal of *Tg (mpx:EGFP)* zebrafish line is detectable in myeloid-specific cells [19]. The *cpa5* gene is a marker gene for mast cells [20]. Meanwhile, mast cells play a pivotal role in the immune response by modulating various cell types, including dendritic cells, macrophages, T cells, B cells, and eosinophils. The *hexb* gene serves as a marker for microglia, which are innate immune cells resident in the central nervous system and are also known as brain macrophages [21]. The *isg15* gene encodes a ubiquitin-like protein that is activated in response to viral infection through IFN - α/β transduction [22]. This activation prompts natural killer (NK) cells and T cells to secrete IFN - γ, a crucial component of the innate immune response that helps combats viral infection. ITK is a member of the Tec family of receptor tyrosine kinase. It plays a pivotal role in regulating the development, differentiation, and function of both conventional and unconventional natural killer T cells. ITK is particularly important in the growth, differentiation, and activation of bone marrow, hypertrophy, and B cells [23]. The *rsad2* gene encodes for an interferon-induced antiviral protein known as viperin (virus inhibitory protein, endoplasmic reticulum-associated, IFN-inducible). This protein has been shown to enhance the production of IFN - β in dendritic cells. It also plays a role in the activation and differentiation of CD4^+^ T cell, and inhibits the secretion of soluble proteins [24].

The main chemical components of Astragalus powder include polysaccharides, flavonoids, and saponins [25]. Specifically, Astragalus polysaccharide (APS), being the most active component in Astragalus powder, has immunomodulatory and anti-tumor effects. APS has been observed to induce macrophage polarization towards M1 phenotype through the Notch signaling pathway. M1 macrophages promote tumor killing response and inhibit tumor growth [26]. In the treatment of immunodeficient models with the immunopotentiator Astragalus powder, we found a significant restorative effect on macrophages across all treatment groups. *Cpa5* and *hexb* were significantly upregulated at a concentration of 1 mg/mL compared to the model group, demonstrating that Astragalus powder possesses a certain immune-enhancing effect (Fig. 3).

10-hydroxydec-2-enoic acid (10 - HDA), a unique component found exclusively in Royal jelly, is known for its potential to promote human health. The enhanced activity of pathways related to DNA/RNA/protein functions mediated by 10 - HDA in the thymus may be crucial for the proliferation, differentiation, and cytotoxicity of T lymphocyte. In the spleen, the induction of pathways involving DNA/RNA/protein activity and stimulation of cell proliferation indicate that these mechanisms are instrumental in the affinity maturation of B lymphocyte, antigen presentation, and macrophage activity [27]. Our results show that treating immunodeficiency models with royal jelly has a significant increase in the population of T cells and macrophages within the thymus. The expression levels of *cap5*, *hexb*, *itk*, and *rads2* in the 200 μg/mL royal jelly-treated group showed no significant deviation from those observed in the control group, thereby proving that royal jelly does indeed have an immune-enhancing effect (Fig. 4).

The basidium, mycelium, and spores of Ganoderma lucidum are rich in bioactive compounds that are known to have various pharmacological effects, including immune regulation [28]. The spore breaking powder of Ganoderma lucidum has been found to promote T cell activation and release of immune factors, thereby exerting tumor immune effects [29]. This powder can also enhance the phagocytic function of macrophages and facilitate the secretion of cytokines [30]. Our research shows that the application of broken Ganoderma lucidum spore powder can significantly increase the content of macrophages and have a certain restoring effect on the expression levels of *cap5*, *hexb*, and *isg15*, proving that Ganoderma lucidum spore powder indeed possesses an immune-boosting effect (Fig. 5).

Bailing Capsules, prepared from Cordyceps sinensis strains through artificial low-temperature fermentation, have been demonstrated by contemporary pharmacological research to enhance cellular immune function. Studies have shown that Cordyceps sinensis can promote the proliferation of T lymphocyte and improve the phagocytic function of mononuclear macrophages [31]. At a concentration of 20 μg/mL, Bailing Capsule has a significant upregulation effect on the content of thymic T cells and myeloid cells. Additionally, the expression levels of *hexb* and *isg15* are higher in the model group compared to the control group, with some results showing no significant difference, thereby proving ther immune-boosting effects of Bailing Capsule (Fig. 6).

In this study, an immunodeficient zebrafish model was established using Tacrolimus and Everolimus to evaluate the impact of four immunopotentiators-Astragalus powder, Royal jelly, Ganoderma lucidum spore powder and Bailing capsule-on immune function and the expression of immune-related genes. The research revealed that all four immunopotentiators were capable of enhancing or restoring the expression of immune cells and immune related genes, although there are certain differences in the ways of enhancement.

## Ethics statement

This study was conducted in accordance with the Guidelines for Care and Use of Laboratory Animals of the Research Ethics Committee of Fuzhou University. The protocol was approved by the Research Ethics Committee of Fuzhou University.

## Author contributions

Conceptualization, Xiaozhen He; methodology, Mei Huang and Zhipeng Jia; software, Mei Huang, Zhipeng Jia and Yiliang Liu; validation, Mei Huang, Zhipeng Jia and Yiliang Liu; formal analysis, Mei Huang, Zhipeng Jia and Yiliang Liu; investigation, Mei Huang, Zhipeng Jia and Xiaozhen He; resources, Xing Lu and Xiaozhen He; data curation, Mei Huang, Zhipeng Jia and Xiaozhen He; writing—original draft preparation, Mei Huang and Zhipeng Jia; writing—review and editing, Mei Huang, Xing Lu and Xiaozhen He; visualization, Mei Huang, Zhipeng Jia and Yiliang Liu; supervision, Xing Lu and Xiaozhen He; project administration, Xing Lu and Xiaozhen He; funding acquisition, Xing Lu and Xiaozhen He. All authors have read and agreed to the published version of the manuscript.

## Funding

This research was funded by Natural Science Foundation of Fujian Province, China (Grant No. 2024J01268), the GDAS’ Project of Science and Technology Development (No. 2022GDASZH-2022010101), National Natural Science Foundation of China (Grant No. 82072986).

## Acknowledgments

We are grateful to the China Zebrafish Resource Center (CZRC) for its zebrafish strains.

## Declaration of competing interest

The authors declare no competing financial interests. One patent application has been filed relating to this work.

## Appendix A. Supplementary data

Supplementary data to this article can be found online at XXX.

### Detailed methods Animal handling

The wild-type (WT) AB strain and the transgenic strains *Tg (rag2: DsRed)*, *Tg (mpx: EGFP)*, and *Tg (lyz: DSRed2)* zebrafish are all obtained from the Chinese Zebrafish Resource Center (CZRC).

The zebrafish work was conducted in accordance with the full animal care and use guidelines, and all procedures were approved by the local institutional animal care committee at the College of Biological Science and Engineering, Fuzhou University.

### Chemical and reagents

Tacolimus (T93035), Everolimus (S43232), and Acridine Orange (S47568) were purchased from Shanghai Yuanye Bio Technology (Shanghai, China, purity ≥ 98%). Lingzhi spore powder was purchased from Jilin Province Tailemei Health Technology Co., Ltd. (Baishan, China), Astragalus powder was purchased from Anhui Risheng Biotechnology Co., Ltd. (Bozhou, China), royal jelly was purchased from Lanxi Jumiyuan Food Technology Co., Ltd. (Lanxi, China), Bailing capsules were purchased from Hangzhou Zhongmei Huadong Pharmaceutical Co., Ltd. (Hangzhou, China), dimethyl sulfoxide was purchased from BBI Life Science Co., Ltd. (Hong Kong, China), and E3 laboratory provided.

### Drug treatments

Healthy zebrafish embryos at 48 hpf were placed into a 12-well plate, with 10 embryos per well in 4 mL of solution (E3). The embryos were then exposed to either a control solution or an immunotherapy inhibitor solution (Tacrolimus + Everolimus) until 120 hpf. The concentrations of the immunotherapy inhibitor for pre-treatment and formal treatment are provided in Supplementary Tables 1 and 2, respectively. The working fluid was replaced with fresh solution every 24 hours to maintain treatment conditions.

The pre-treatment and formal treatment concentrations of immune enhancers such as Astragalus powder, Royal jelly, Ganoderma lucidum spore powder and Bailing capsules are shown in Supplementary Tables 3 and 4, respectively.

### Acridine orange (AO) staining

WT zebrafish at 48 hpf were exposed to a control solution (E3) and an immune combination inhibitor (Tacrolimus + Everolimus) solution until for 72 h. At 120 hpf, zebrafish treated with wild-type and immune combination inhibitors were stained with acridine orange. The procedure began by washing the zebrafish with clean water. Subsequently, the culture water was replaced with 5 μg/mL acridine orange PBS solution, and the zebrafish were placed in a 28 [incubator in the dark for 30 minutes. After washing with PBS solution, fish were placed on a shaker for 5 minutes and a process that was repeated three times.

### *In vivo* imaging of zebrafish embryos

To investigate the total number, distribution, and recruitment of immune cells, 120 hpf transgenic fish larvae from both control and treatment groups were anesthetized with 0.16 mg/mL tricaine in embryo culture medium. Images of nose, tail or the entire body, depending on the research model were captured using a Nikon SMZ188 fluorescence stereomicroscope. The number of fluorescent/stained cells was counted visually, and the fluorescence intensity for various parameters (apoptotic cells, thymus T cells, myeloid cells, macrophages) within the region of interest (ROI) was analyzed using ImageJ.

### Quantitative real-time PCR validation

Real-time fluorescence quantitative PCR (RT-qPCR) was used to detect the gene transcript levels of five immune-related genes (*itk*, *cap5*, *hexb*, *isg15* and *rsad2*). The cDNAs of all samples were used as templates for PCR. Specific primers were designed using Primer premier 5 software based on the coding DNA (cDNA) sequence of each gene retrieved on Ensemble Genome Browser (http://asia.ensembl.org/index.html) (Supplementary Table 5). The reaction volume was 20 μL, in which 2 × chamQ Universal SYBR qPCR Master Mix mixture was 5 μL, ddH2O 10.5 μL, template 2.5 μL, and primer 1 μL. Cycling conditions were 95 for 3 min, 94 for 10 s, 60 for 30 s, 95 for 15 s. The cycle was performed 40 times, and 3 parallel experiments were performed to verify the results. The results were calculated by 2^-ΔΔCT^ analysis method.

### Statistical analyses

Statistical analysis was performed using Prism 8.0 (GraphPad Software, CA, USA). The sample size was not predetermined; instead, the experiment was repeated three times to ensure robustness. Embryos were randomly assigned to each experimental condition and analyzed in a blinded sample. No data were excluded from the analysis. The data is presented as mean ± SEM. Statistical comparisons between the two groups were performed using the independent two-sample *t*-test.Data from multiple groups were analyzed using a one-way analysis of variance (ANOVA) and Tukey’s multiple range test. A value of *P* < 0.05 is considered statistically significant.

## Data availability

Data will be made available on request.

## Notes

### Competing Interest Statement

The authors have declared no competing interest.

### Summary of Updates

In this revised version, we have included Jun Zhou as a co-author based on their substantial contributions to data updating and independent verification. Furthermore, all relevant data points are now presented as scatter plots to enable direct visual assessment of data distributions and relationships by readers.

## References

[1] A.M. van der Sar, R.J.P. Musters, F.J.M. van Eeden, B.J. Appelmelk, C. Vandenbroucke-Grauls, W. Bitter, Zebrafish embryos as a model host for the real time analysis of Salmonella typhimurium infections, Cellular Microbiology 5(9) (2003) 601–611, 10.1046/j.1462-5822.2003.00303.x

[2] Y. Li, Z. Jia, S. Zhang, X. He, Progress in Gene-Editing Technology of Zebrafish, Biomolecules 11(9) (2021), 10.3390/biom11091300.

[3] G.J. Lieschke, A.C. Oates, M.O. Crowhurst, A.C. Ward, J.E. Layton, Morphologic and functional characterization of granulocytes and macrophages in embryonic and adult zebrafish, BLOOD 98(10) (2001) 3087–3096, 10.1182/blood.V98.10.3087.

[4] S.H. Lam, H.L. Chua, Z. Gong, T.J. Lam, Y.M. Sin, Development and maturation of the immune system in zebrafish, Danio rerio: a gene expression profiling, in situ hybridization and immunological study, Dev Comp Immunol 28(1) (2004) 9–28, 10.1016/S0145-305X(03)00103-4.

[5] N.S. Trede, D.M. Langenau, D. Traver, A.T. Look, L.I. Zon, The use of zebrafish to understand immunity, Immunity 20(4) (2004) 367–379, 10.1016/S1074-7613(04)00084-6.

[6] J.J. Perez-Villar, G.S. Whitney, M.T. Sitnick, R.J. Dunn, S. Venkatesan, K. O’Day, G.L. Schieven, T.A. Lin, S.B. Kanner, Phosphorylation of the linker for activation of T-cells by Itk promotes recruitment of Vav, Biochemistry 41(34) (2002) 10732–10740, 10.1021/bi025554o.

[7] S.A. Renshaw, C.A. Loynes, D.M. Trushell, S. Elworthy, P.W. Ingham, M.K. Whyte, A transgenic zebrafish model of neutrophilic inflammation, Blood 108(13) (2006) 3976–3978, 10.1182/blood-2006-05-024075.

[8] M.J. Redd, G. Kelly, G. Dunn, M. Way, P. Martin, Imaging macrophage chemotaxis in vivo: studies of microtubule function in zebrafish wound inflammation, Cell Motil Cytoskeleton 63(7) (2006) 415–422, 10.1002/cm.20133.

[9] C. Hall, M.V. Flores, T. Storm, K. Crosier, P. Crosier, The zebrafish lysozyme C promoter drives myeloid-specific expression in transgenic fish, BMC Dev Biol 7 (2007), 42. 10.1186/1471-213X-7-42.

[10] N.D. Meeker, N.S. Trede, Immunology and zebrafish: Spawning new models of human disease, Developmental and Comparative Immunology 32(7) (2008) 745–757, 10.1016/j.dci.2007.11.011.

[11] J.J. Fung, Tacrolimus and transplantation: A decade in review, TRANSPLANTATION 77(9) (2004) S41–S43, 10.1097/01.TP.0000126926.61434.A5.

[12] S.C. Ong, R.S. Gaston, Thirty Years of Tacrolimus in Clinical Practice, Transplantation 105(3) (2021) 484–495, 10.1097/TP.0000000000003350.

[13] Zhang. M.S, Sun. T, 2010 Everolimus, Chinese Journal of Medicinal Chemistry 20 (06), 553-554. 10.14142/j.cnki.cn21-1313/r.2010.06.018.

[14] L. Gullestad, S.A. Mortensen, H. Eiskjoer, G.C. Riise, L. Mared, O. Bjortuft, B. Ekmehag, K. Jansson, S. Simonsen, E. Gude, B. Rundqvist, H.E. Fagertun, D. Solbu, M. Iversen, Two-Year Outcomes in Thoracic Transplant Recipients After Conversion to Everolimus With Reduced Calcineurin Inhibitor Within a Multicenter, Open-Label, Randomized Trial, Transplantation 90(12) (2010) 1581–1589, 10.1097/TP.0b013e3181fd01b7.

[15] S.S. Wang, N.K. Chou, N.H. Chi, S.C. Huang, I.H. Wu, C.H. Wang, H.Y. Yu, Y.S. Chen, C.I. Tsao, W.J. Ko, C.T. Shun, Clinical Experience of Tacrolimus With Everolimus in Heart Transplantation, Transplantation Proceedings 44(4) (2012) 907–909, 10.1016/j.transproceed.2012.01.094.

[16] P. De Simone, F. Nevens, L. De Carlis, H.J. Metselaar, S. Beckebaum, F. Saliba, S. Jonas, D. Sudan, J. Fung, L. Fischer, C. Duvoux, K.D. Chavin, B. Koneru, M.A. Huang, W.C. Chapman, D. Foltys, S. Witte, H. Jiang, J.M. Hexham, G. Junge, Everolimus With Reduced Tacrolimus Improves Renal Function in De Novo Liver Transplant Recipients: A Randomized Controlled Trial, American Journal of Transplantation 12(11) (2012) 3008–3020, 10.1111/j.1600-6143.2012.04212.x.

[17] N.A. Clipstone, G.R. Crabtree, Calcineurin is a key signaling enzyme in T lymphocyte activation and the target of the immunosuppressive drugs cyclosporin A and FK506, Ann N Y Acad Sci 696 (1993) 20–30, 10.1038/357695a0.

[18] U. Saran, M. Foti, J.F. Dufour, Cellular and molecular effects of the mTOR inhibitor everolimus, Clin Sci (Lond) 129(10) (2015) 895–914, 10.1042/CS20150149.

[19] H. Yuan, J. Zhou, M. Deng, Y. Zhang, Y. Chen, Y. Jin, J. Zhu, S.J. Chen, H. de The, Z. Chen, T.X. Liu, J. Zhu, Sumoylation of CCAAT/enhancer-binding protein α promotes the biased primitive hematopoiesis of zebrafish, Blood 117(26) (2011) 7014–7020, 10.1182/blood-2010-12-325712.

[20] O. Goldmann, T. Sauerwein, G. Molinari, M. Rohde, K.U. Förstner, E. Medina, Cytosolic Sensing of Intracellular Staphylococcus aureus by Mast Cells Elicits a Type I IFN Response That Enhances Cell-Autonomous Immunity, J Immunol 208(7) (2022) 1675–1685, 10.4049/jimmunol.2100622.

[21] U.K. Hanisch, H. Kettenmann, Microglia: active sensor and versatile effector cells in the normal and pathologic brain, Nat Neurosci 10(11) (2007) 1387–1394, 10.1038/nn1997.

[22] P.F. Dos Santos, D.S. Mansur, Beyond ISGlylation: Functions of Free Intracellular and Extracellular ISG15, J Interferon Cytokine Res 37(6) (2017) 246–253, 10.1089/jir.2016.0103.

[23] J.J. Perez-Villar, G.S. Whitney, M.T. Sitnick, R.J. Dunn, S. Venkatesan, K. O’Day, G.L. Schieven, T.A. Lin, S.B. Kanner, Phosphorylation of the linker for activation of T-cells by Itk promotes recruitment of Vav, Biochemistry 41(34) (2002) 10732–10740, 10.1021/bi025554o.

[24] A.B. Dumbrepatil, S. Ghosh, K.A. Zegalia, P.A. Malec, J.D. Hoff, R.T. Kennedy, E.N.G. Marsh, Viperin interacts with the kinase IRAK1 and the E3 ubiquitin ligase TRAF6, coupling innate immune signaling to antiviral ribonucleotide synthesis, Journal of Biological Chemistry 294(17) (2019) 6888–6898, 10.1074/jbc.RA119.007719.

[25] Ma, Y., Hu, J., Wu, W., Duan, Y., Fan, C.C., Feng, T., Wang, X., Wu, X.H., 2022 Research progress on chemical components and pharmacological effects of Astragalus membranaceus, Acta Chinese Medicine and Pharmacology 50 (04), 92–95, 10.19664/j.cnki.1002-2392.220092.

[26] W. Wei, Z.P. Li, Z.X. Bian, Q.B. Han, Astragalus Polysaccharide RAP Induces Macrophage Phenotype Polarization to M1 via the Notch Signaling Pathway, Molecules 24(10) (2019), 10.3390/molecules24102016.

[27] P. Fan, B. Han, H. Hu, Q. Wei, X. Zhang, L. Meng, J. Nie, X. Tang, X. Tian, L. Zhang, L. Wang, J. Li, Proteome of thymus and spleen reveals that 10-hydroxydec-2-enoic acid could enhance immunity in mice, Expert Opin Ther Targets 24(3) (2020) 267–279, 10.1080/14728222.2020.1733529.

[28] B.S. Sanodiya, G.S. Thakur, R.K. Baghel, G.B. Prasad, P.S. Bisen, Ganoderma lucidum: a potent pharmacological macrofungus, Curr Pharm Biotechnol 10(8) (2009) 717–742, 10.2174/138920109789978757.

[29] S. Cheng, D. Sliva, Ganoderma lucidum for cancer treatment: we are close but still not there, Integr Cancer Ther 14(3) (2015) 249–257, 10.1177/1534735414568721.

[30] Wang. Y, Jin. J, Jin. Z, Li. J, Mao. P.J, Yang. Yi., Huang, J.J, Li. M, 2016 Research on Enhancing Immune Function with Ganoderma Spore Powder, Chinese Modern Applied Pharmacy 33 (05) 544–546, 10.13748/j.cnki.issn1007-7693.2016.05.006.

[31] Chen. Di, Wu. F, 2017. Clinical effects and the impact of the organism cellular immunity function and micro-inflammation state in primary nephroticsyndrome treating with Bailing capsule, Shanxi Traditional Chinese Medicine 38 (12), 1670–1672, 10.3969/j.issn.1000-7369.2017.12.017.

