## Supplemental Table 1-2 for "A zebrafish immunodeficiency model induced by the combination of Tacrolimus and Everolimus for rapid assessment of immune-enhancing agents"

Supplemental Table 1 Pre-treatment concentration

| | Control | Pre-G1 | Pre-G2 | Pre-G3 | Pre-G4 |
|---|---|---|---|---|---|
| Tacrollimus（ng/mL） | 0 | 0.1 | 1 | 5 | 10 |
| Everolimus（μg/mL） | 0 | 1 | 15 | 30 | 45 |

Supplemental Table 2 Final drug concentration ratio

| | Control | G1 | G2 | G3 | G4 |
|---|---|---|---|---|---|
| Tacrollimus（ng/mL） | 0 | 0.5 | 1.5 | 2.5 | 3.5 |
| Everolimus（μg/mL） | 0 | 5 | 10 | 15 | 20 |

Supplemental Table 3 Pre-experimental concentrations of immunomodulators

| Astragalus powder (mg/mL) | Ganoderma lucidum spore powder(μg/mL) | Royal jelly (μg/mL) | Bailing capsule (μg/mL) |
|---|---|---|---|
| 0 | 0 | 0 | 0 |
| 1 | 100 | 100 | 50 |
| 2 | 500 | 200 | 100 |
| 3 | 1000 | 300 | 150 |
| 4 | 1500 | 400 | 200 |
| 5 | 2000 | 500 | 250 |

Supplemental Table 4 Concentration in the experimental group of immunomodulators

| Astragalus powder (mg/mL) | Ganoderma lucidum spore powder(μg/mL) | Royal jelly (μg/mL) | Bailing capsule (μg/mL) |
|---|---|---|---|
| 1 | 100 | 100 | 20 |
| 2 | 500 | 200 | 50 |
| 3 | 1000 | 300 | 100 |

Supplemental Table 5 Primer sequences used for qRT-PCR.

| Gene symbol (gene ID) | Primer Sequence(5'-3' ) |
|---|---|
| *isg15*-RT-F | TAATGCCACAGTCGGTGAA |
| *isg15*-RT-R | AGGTCCAGTGTTAGTGATGAGC |
| *rsad2*-RT-F | GCAAAGCGAGGGTTACGAC |
| *rsad2*-RT-R | CTGCCATTACTAACGATGCTGAC |
| *cpa5*-RT-F | GGGCGAACATCTTGGCCTTA |
| *cpa5*-RT-R | CACGTTGCTCATCATCCAACA |
| *hexb*--RT-F | TGAGCCGTACTTTCCCAGAG |
| *hexb*--RT-R | GCAGATCCTTTATGCCATTGCC |
| *itk*-RT-F | CGGTTGCCTTACGGAGTTCT |
| *itk*-RT-R | AACAGTTTCTCGCAGCCAAG |
| *β-actin*-F | CGAGCAGGAGATGGGAACC |
| *β-actin*-R | CAACGGAAACGCTCATTGC |

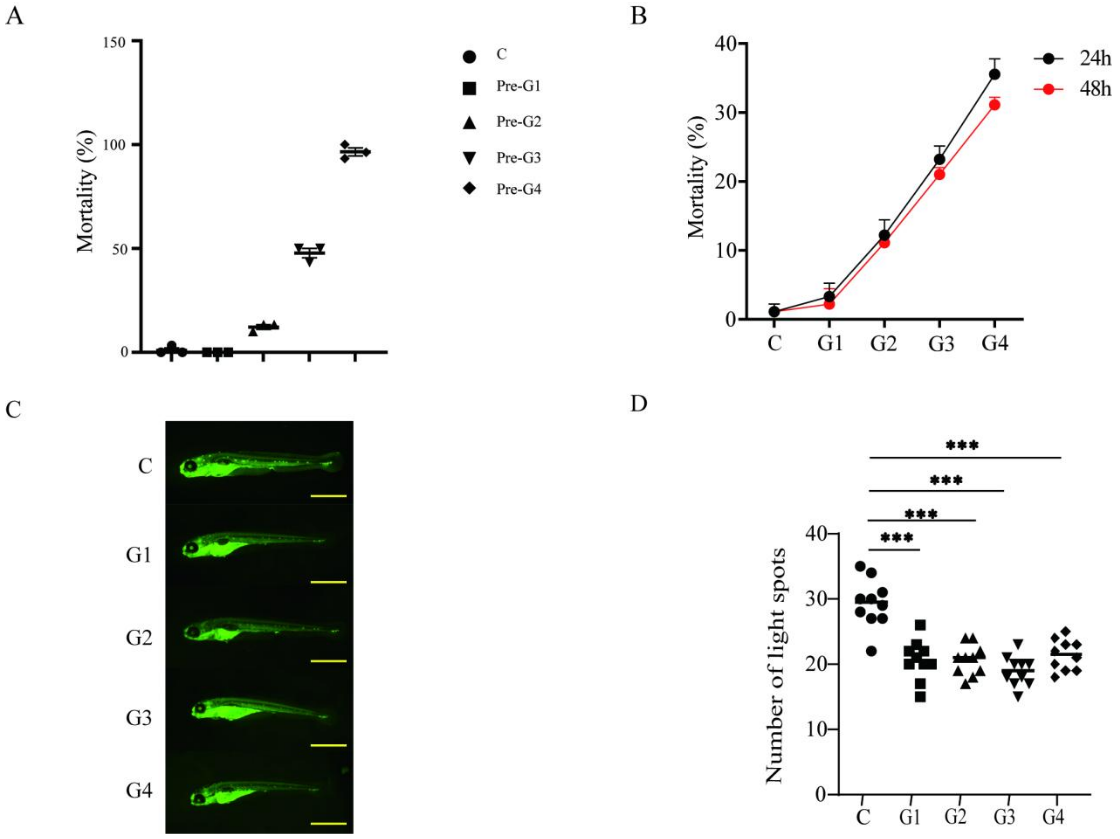


**Supplementary Figure 1.** Acquisition of immunodeficiency model. (A) Concentration ratio pre experiment: 120 hpf mortality rate. (B) Shows the mortality rate of zebrafish in the Control group with different ratios of tacrolimus and everolimus in G1, G2, G3, and G4. (C) Control group, G1, G2, G3 and G4 120 hpf zebrafish acridine orange stained pictures. (D) Statistics of the number of apoptotic light spots in Control group, G1, G2, G3 and G4 120 hpf zebrafish acridine orange stained.

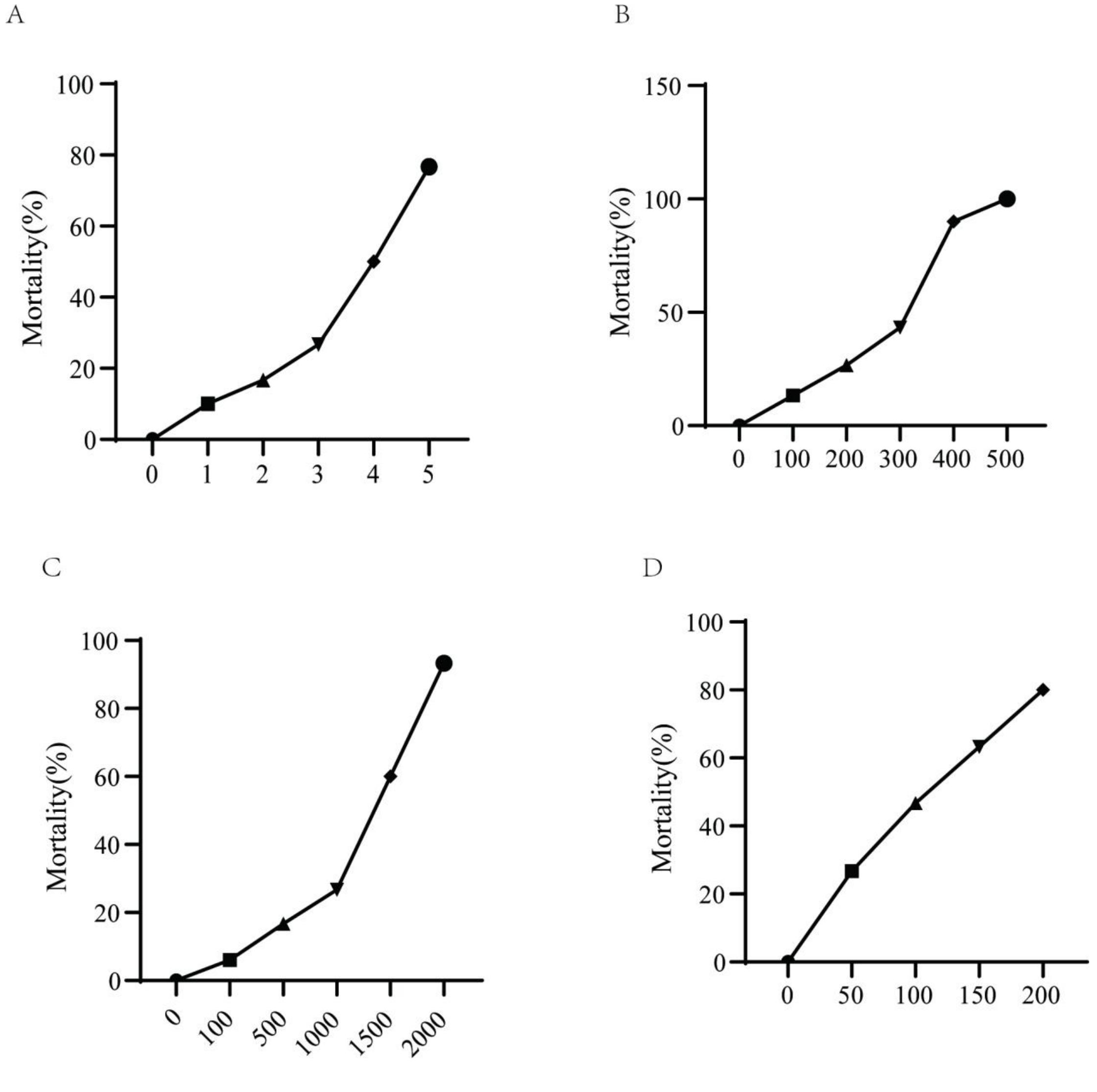


**Supplementary Figure 2.** The effects of immune modulators powder on the zebrafish were found to be concentration-dependent, with increased immune enhancement observed as the drug concentration increased. (A) 0, 1, 2, 3, 4 and 5 mg/mL Astragalus powder 48 hpf - 120 hpf treatment for zebrafish mortality rate. (B) 0, 100, 200, 300, 400 and 500 μg/mL Royal jelly 48 hpf - 120 hpf treatment for zebrafish mortality rate. (C) 0, 100, 500, 1000, 1500 and 2000 μg/mLGanoderma lucidum spore powder 48 hpf - 120 hpf treatment for zebrafish mortality rate. (D) The mortality rate of zebrafish treated with 0, 50, 100, 150 and 200 μg/mL Bailing capsules at 48 hpf-120 hpf.
